# Biomimetic Aorta-Gonad-Mesonephros-on-a-Chip to Study Human Developmental Hematopoiesis

**DOI:** 10.1101/837856

**Authors:** Ryohichi Sugimura, Ryo Ohta, Chihiro Mori, Alina Li, Takafumi Mano, Emi Sano, Kaori Kosugi, Tatsutoshi Nakahata, Akira Niwa, Megumu K. Saito, Yu-suke Torisawa

## Abstract

We established a multilayer microfluidic Aorta-Gonad-Mesonephros (AGM)-on-a-chip to emulate developmental hematopoiesis from human pluripotent stem cells. We show that the AGM-chip efficiently derives endothelial-to-hematopoietic transition (EHT) in the presence of mesenchymal stroma and endothelial cells. The AGM-chip could dissect the cellular and molecular mechanisms of human developmental hematopoiesis.

## Introduction

The hematopoietic system sustains life through an almost limitless production of blood cells and resistance to infection. The generation of all blood cells originates in the embryonic tissue called aorta-gonad-mesonephros (AGM)^1,2^. At around 12-15 days post coitum (dpc) of human embryos, a subset of endothelial cells called hemogenic endothelium (HE) in the AGM are specified to hematopoietic progenitor cells (HPCs) through endothelial-to-hematopoietic transition (EHT)^2^. Capturing the precise mechanisms of human hematopoietic development, especially EHT process, will benefit efforts to generate blood cells *in* vitro for cell therapy, drug screening and disease modeling^3–5^. Fluidic systems have been utilized to model the environment of HE^6,7^; however, HE cells are typically monocultured on a plate without feeder cells. Herein we propose a novel method to model human hematopoietic development using the organ-on-a-chip technology^8–12^. We developed a microfluidic AGM-on-a-chip which resembles key features of the cellular components of human AGM^2^, including the HE, mesenchymal stroma (MS) and endothelial cells (ECs). We previously isolated a MS cell line from murine AGM that supports EHT *in vitro* (AGMS-3)^13^, which we use in the current study to recapitulate mesenchymal stroma in the AGM. We also used human umbilical vein endothelial cells (HUVECs) as a platform of ECs because they have the capacity to support hematopoiesis *in vitro*^14^ and are easy to handle. We demonstrate that the AGM-chip efficiently induces EHT from human induced pluripotent stem cells (hiPSCs) compared with regular suspension culture, providing a robust generation of human blood cells. Of note, the presence of MS and ECs renders EHT with functional HPCs *in vitro*. The perfusion culture ensured recovery of HPCs from the device, allowing for connection to other types of organs-on-chips. We propose our AGM-chip as a novel platform to study the cellular and molecular mechanisms of human hematopoietic development.

## Results and discussion

The AGM-chip is biomimetic both in terms of structure and cellular interactions during hematopoietic development (Fig. 1A and 1B). We employed hiPSC-derived hemogenic endothelium (hiPSC-HE), HUVECs as a model of ECs, and AGMS-3 cells as a model of MS into the AGM-chip. The device consists of two polydimethylsiloxane (PDMS) microchannels separated by a 3-um porous membrane^15^, which allows cellular interactions between each cell layer^8^ (Fig. 1B and 1C). This system realizes to recover differentiated cells from the outlet for analyses by flowing fluid through the microchannel. We inputted hiPSC-HE and AGMS-3 cells in the upper layer, and HUVECs in the lower layer (Fig. 1B). We coated the membrane with fibronectin and inputted AGMS-3 cells and HUVECs in the device prior to the injection of hiPSC-HE cells (Fig. 1D). Hematopoietic specification of hiPSC-HE was verified on the chip in which hiPSC-HE cells were cultured directly onto a layer of AGMS-3 cells and a layer of HUVECs formed underneath (Fig. 2). We examined the hematopoietic specification of GFP-expressing hiPSC-HE cells in the AGM-chip. We observed the formation of round-shaped GFP+ cells after 6 days in culture on-chip, indicating hematopoietic transition. The observation is consistent with the notion that ECs and MS coax hematopoietic specification of HE^16^ (Fig. 2).

**Fig. 1.**
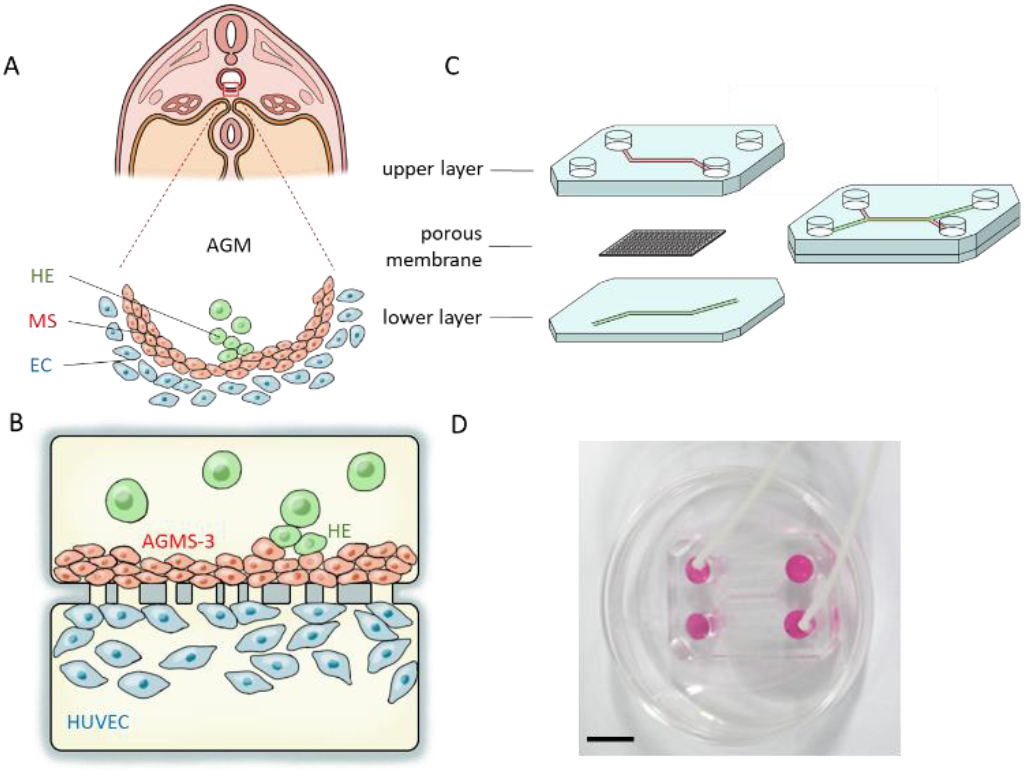
Construction of Aorta-Gonad-Mesonephros (AGM)-on-a-chip. (A) Scheme of the AGM region in the human embryo. Hemogenic endothelium (HE), mesenchymal stroma (MS) and endothelial cells (EC) compose the AGM. (B) Scheme of the AGM-chip. HE and AGMS-3 cells are placed in the upper layer and HUVECs in the lower layer. A porous membrane (3 µm pores) in the middle allows the cells to interact through the layers. (C) Structure of the AGM-chip. The upper and lower layers are made of PDMS and sandwich the porous membrane. Culture medium is infused from a reservoir (6 mm in diameter) and perfused through a microchannel using a micro peristaltic pump. (D) Photograph of the AGM-chip. Scale bar = 10 mm.

**Fig. 2.**
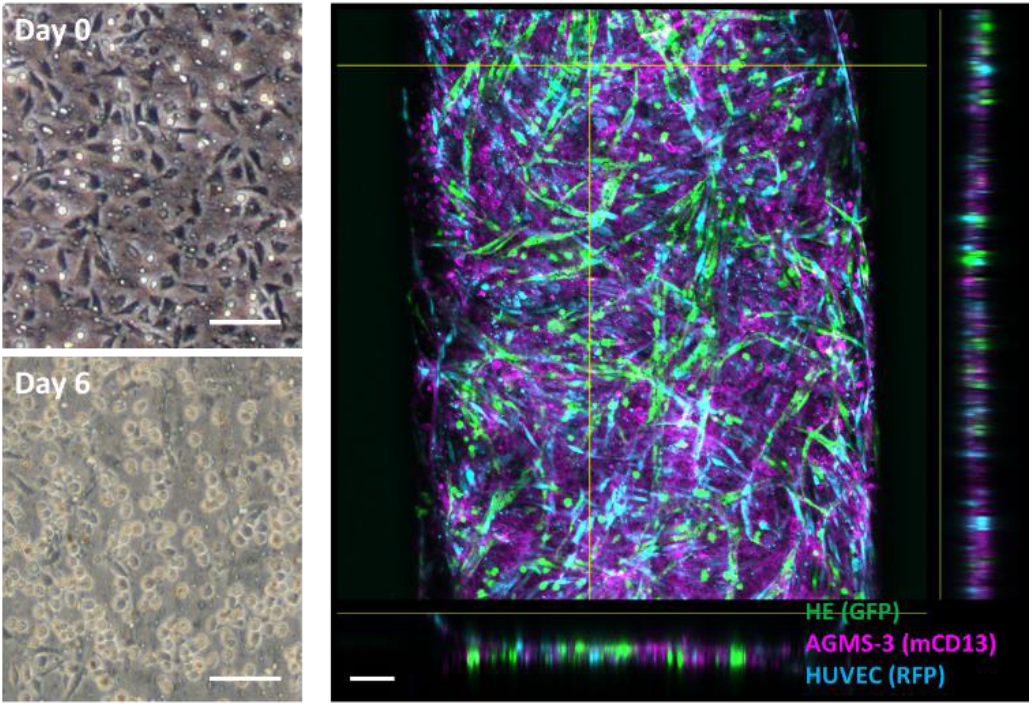
Cellular components of AGM-chip. Left; A bright-field microscopy image of HE seeded on the chip without feeder cells. The formation of round-shaped hematopoietic cells is shown. Right; A confocal fluorescence micrograph of the AGM-chip with feeder cells consisting of GFP-expressing HE (green), mCD13-stained AGMS-3 cells (magenta), and RFP-expressing HUVECs (cyan). Cross-sectional images are shown on the right and bottom. Scale bar = 100 µm.

We then examined EHT as the yields of hematopoietic progenitor cells (HPCs) from an entire AGM-chip (Fig. 3A). We found that both the AGM-chip and suspension culture produced HPCs at slimier yields at both 1- and 2-weeks of culture. The presence of feeder cells (AGMS-3 cells and HUVECs) significantly rendered HPCs from the AGM-chip at 2 weeks (with and without feeder, p=0.013) (Fig. 3B). Consistent with the number of HPCs, colony-forming-unit (CFU) capacity depends on the presence of feeder cells at 2-weeks of culture (Fig. 3C), indicating for the role of feeder cells in sustaining functional HPCs in culture. Moreover, CFU capacity of the AGM-chip was relatively higher than that of suspension culture at both 1-week (p=0.05) and 2-weeks (p=0.08) of culture in the presence of feeder cells (Fig. 3C). These data demonstrate the advantage of our AGM-chip in producing HPCs over suspension culture, indicating microenvironmental cues play an important role in the induction of EHT. To assess the production of HPCs from the AGM-chip, we harvested cells from the outflow of the device by perfusing culture medium using a micro peristaltic pump at a flow rate of 20 µL min^−1^ (Fig. 4A). We found that around 47% of the cells in the outflow were CD34+CD45+ HPCs and confirmed their colony-forming capacity (Fig. 4B and 4C). These data demonstrate that the AGM-chip elicits HPCs for prospective connection to other types of organs-on-chips.

**Fig. 3.**
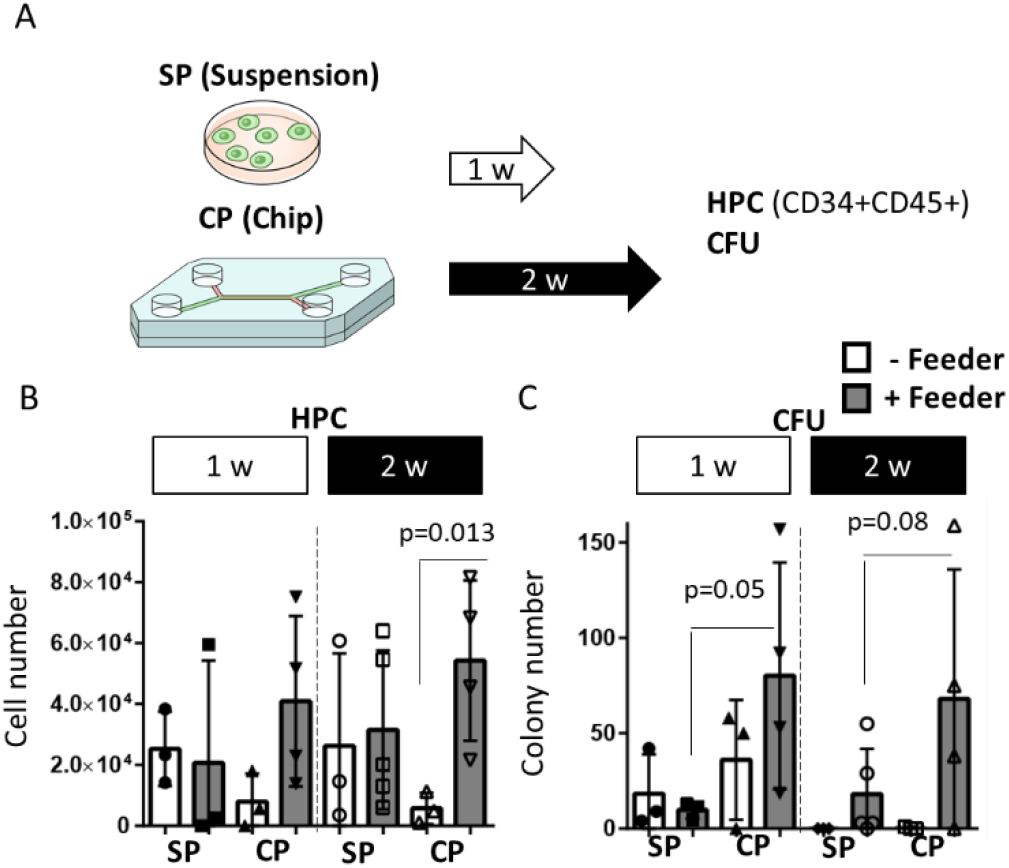
Fig. 3 EHT in the AGM-chip. (A) Experimental scheme. HE cells were seeded with or without feeder cells (AGMS-3 and HUVEC). Aftetr 1 or 2 weeks in culture, whole cells from the chip were collected and analyzed with HPC markers (CD34 and CD45) and with methylcellulose hematopoietic CFU assay. The bar graphs show the number of HPCs (B) (1 w; n=3, 2 w; n=3-5) and CFU (C) (1 w; n=3, 2 w; n=3-5). Each dot represents the result from one biologically independent experiment.

**Fig. 4.**
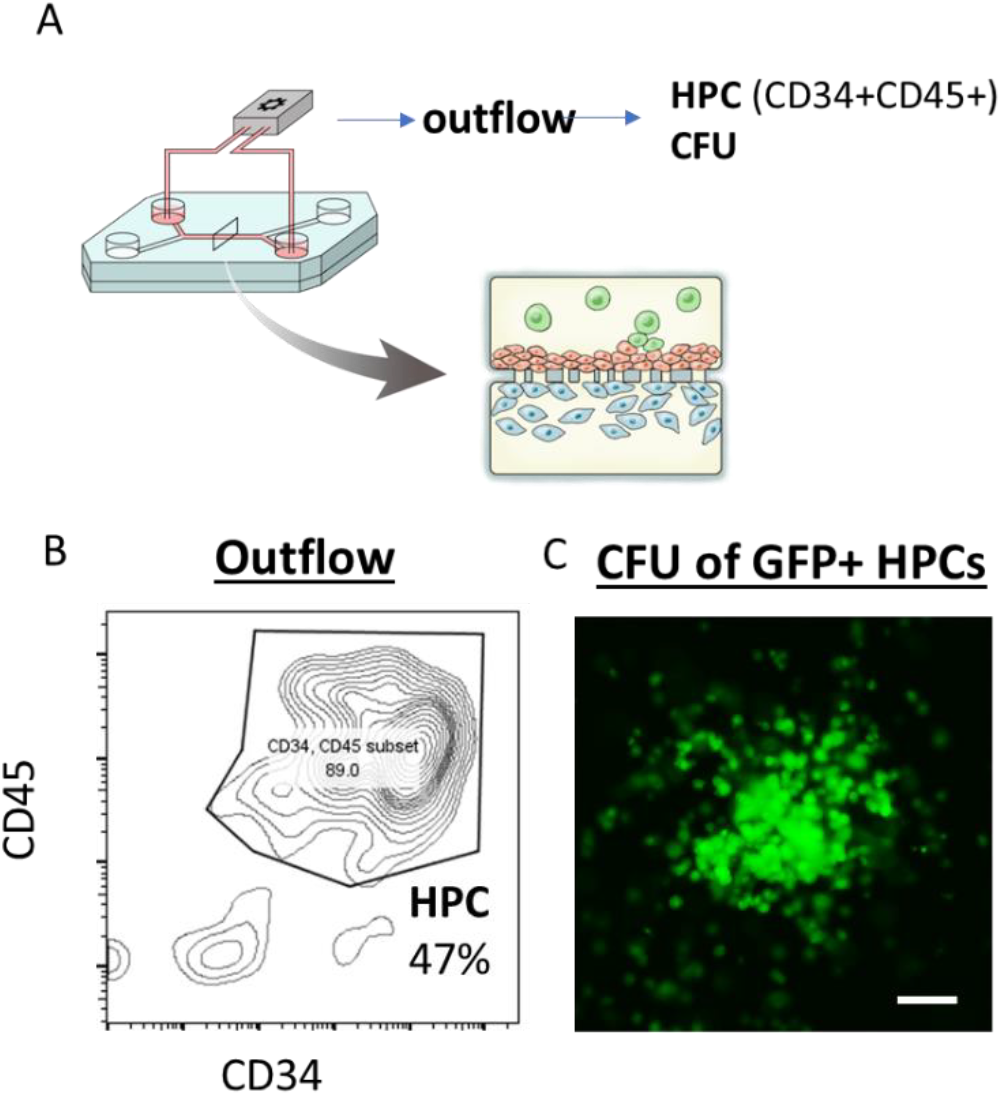
Collection of HPCs from the outflow of AGM-chip. (A) Experimental scheme. GFP-labeled HE, AGMS-3 and HUVEC are seeded. After 1-week culture, cells from the outflow were analyzed with the HPC markers CD34 and CD45 and with CFU. (B) FACS plot of the outflow cells with CD34 and CD45. Average percentage of HPC was shown from n=3. (C) A GFP+ colony under fluorescence microscopy. Scale bar = 200 µm

## Conclusions

In this paper, we present an *in vitro* AGM-on-a-chip platform as biomimetic AGM which is where blood cells emerge during development^17^. We demonstrated that culture in the AGM-chip yields more EHT than suspension cell culture indicated by the presence of HPCs capable of colony formation *in vitro*. Furthermore, we demonstrated that feeder cells support EHT. We expect the AGM-chip could be a powerful experimental tool with which end users can assess the effects of target signaling, small molecules, genetic interventions in human developmental hematopoiesis and connect with other types of organs-on-chips.

## Materials and methods

### Cell lines

All the experiments of this study were performed with 409B2 iPSCs or CBA11 iPSCs^18^. HUVECs were either purchased from Angio-Proteomie (GFP-HUVEC, cAP-01001GFP) or Lonza (issue Acquisition Number: 29000). AGMS-3 was isolated and cultured as described previously^13^.

### Cell culture

The maintenance of hiPSCs was done using iMatrix-511 (Matrixome) in mTeSR1 medium (STEMCELL Technologies). Culture medium was changed every other day, and the cells were passaged as single cells every 7 days using TrypLE Express (Life technologies). HUVECs were cultured in endothelial cell growth medium-2 (EGM-2, Lonza) with 100 U mL^−1^ penicillin and 100 U mL^−1^ streptomycin (Nacalai Tesque). All the experiments were conducted using HUVECs of passage 7 or lower. AGMS-3 cells were cultured in alpha-MEM (Thermo Fischer Scientific) containing 15% fetal bovine serum (F7524, Sigma-Aldrich), 100 U mL^−1^ penicillin, and 100 U mL^−1^ streptomycin.

### Differentiation of hemogenic endothelial cells

hiPSC spheroids were formed as described previously^18^, suspended in mTeSR1 medium containing 1.25 µg mL^−1^ iMatrix-511, and plated onto non-coated 100mm culture plates (Corning). After three days in culture, the medium was replaced with Essential 8 (Life technologies) containing 4 µM CHIR99021 (WAKO), 80 ng mL^−1^ BMP4 (R&D systems), and 80 ng mL^−1^ VEGF (R&D systems). After two more days in culture, the medium was replaced with differentiation medium consisting of Essential 6 with 1 µM SB431542, 80 ng mL^−1^ VEGF, and 100 ng mL^−1^ Stem cell factor (SCF, R&D). Finally, two days later, the cells were harvested with TrypLE express, and CD34+ cells were isolated by MACS (CD34 microbeads, (#130-046-703 Miltenyi Biotec). Cytokines were purchased from R&D otherwise indicated.

### Colony forming unit assay

After 1 or 2 weeks in culture, CD45+ cells were sorted by MACS (#130-045-801, Miltenyi Biotec) and cultured at a concentration of 1×10^3^ cells mL^−1^ in one well of a 6-well plate (#1008; Becton-Dickinson) with 1 mL MethoCult GF+ semisolid medium (#4435, STEMCELL Technologies). Colonies were counted after 14 days of incubation using an inverted microscope by three individuals.

### Flow Cytometry

Cells were stained with 1:20 dilution of each antibody for at least 30 min on ice in the dark with the antibodies CD34-PE (#304012, Biolegend) and CD45-APC (#343506, Biolegend). Acquisitions were done on a BD FACSAria II cell sorter or BD LSRFortessa cytometer. Flow cytometry data were analyzed using FlowJo V.10.

### Fabrication of Microfluidic Device

The microfluidic device consists of two layers of microchannels separated by a semipermeable membrane. The microchannel layers were fabricated from poly(dimethylsiloxane) (PDMS) using a soft lithographic method^19^. PDMS prepolymer (Sylgard 184, Dow Corning) at a ratio of 10:1 base to curing agent was cast against a mold composed of SU-8 2150 (MicroChem) patterns formed on a silicon wafer. The cross-sectional size of the microchannels was 1 mm in width and 250 µm in height. To introduce solutions into the channels, access holes were punched through the PDMS using a 6-mm biopsy punch (Kai Corporation). Two PDMS layers were bonded to a semipermeable PET membrane containing 3.0 µm pores (#353091, Falcon) using a thin layer of liquid PDMS prepolymer as a mortar^15^. PDMS prepolymer was spin-coated (4000 rpm for 60 s) onto a glass slide. Subsequently, both the top and bottom channel layers were placed on the glass slide to transfer the thin layer of PDMS prepolymer onto the embossed PDMS surfaces. The membrane was then placed onto the bottom layer and sandwiched with the top layer. The combined layers were left at room temperature for 1 day to remove air bubbles and then put into an oven at 60 °C overnight to cure the PDMS glue.

### Microfluidic Cell Culture

The microfluidic devices were sterilized by placing under UV light for 2 hours. Following the sterilization, the semipermeable membrane of the device was coated with fibronectin (#33016015, Gibco) by incubating the microchannels with a fibronectin solution (100 µg mL^−1^ in PBS) at room temperature for 2 hours. The channels were then rinsed with culture medium prior to cell seeding. A suspension of HUVECs (5 × 10^4^ cells per 10 µL) was introduced into the bottom channel, and then the device was inverted and incubated at 37 °C for 1 hour to allow the cells to adhere onto the bottom side of the membrane. Once cellular attachment was confirmed, the device was flipped back, and the microchannels were filled with culture medium (EGM-2). After 1 day in culture, a suspension of AGMS-3 cells (2 × 10^4^ cells per 10 µL) was introduced into the top channel. After incubation at 37 °C for 1 hour, culture medium (alpha-MEM) was introduced into the top channel and incubated at 37 °C for 1 day to form layers of endothelium and mesenchymal stroma. Subsequently, a suspension of HE cells (2.5 × 10^4^ cells per 5 µL) was introduced into the top channel and incubated at 37 °C for 2 hours to enable cell attachment. The reservoirs were filled with EHT medium (Stemline II Hematopoietic Stem Cell Expansion Medium S0192-500 Sigma-Aldrich) supplemented with 50 ng mL^−1^ SCF, 20 ng mL^−1^ thrombopoietin (TPO), 50 ng mL^−1^ Fms-related tyrosine kinase 3 ligand (Flt-3L), 20 ng mL^−1^ Interleukin-6/Interleukin-6 receptor alpha (IL-6/IL-6Rα), insulin-transferrin-selenium-ethanolamine, L-glutamine, and Penicillin-Streptomycin-Amphotericin B (150 µL in each reservoir), and the device was put in an incubator (37 °C, 5% CO^2^). The devices were cultured for up to 14 days with replacement of the culture medium every 2 or 3 days. To harvest cells from the AGM chip, the outflow from the device was collected by perfusing culture medium using a micro peristaltic pump (RP-TXP5F, Aquatech Co., Ltd.) at a flow rate of 20 µL min^−1^ (0.25 dyne cm^−2^) 6 hours after seeding HE cells.

### Suspension Culture

To form a feeder layer, HUVECs and AGMS-3 cells were cultured in a 12 well plate (TPP). 5 × 10^4^ HUVECs were seeded into a plate and cultured in EGM-2. After 1 day in culture, 5 × 10^4^ AGMS-3 cells were seeded into the plate and cultured in 50% EGM-2 and 50% alpha-MEM for 1 day. Subsequently, a suspension of HE cells (2.5 × 10^4^ cells) was seeded onto the plate and cultured in EHT medium (2 mL). Culture medium was replaced once a week.

### Immunofluorescence

Cells were fixed with 4% paraformaldehyde (PFA, Nacalai Tesque) for 20 min, washed with PBS, and permeabilized with 0.1% Triton X-100 (Sigma-Aldrich) for 15 min. The cells were then incubated with blocking buffer containing 1% bovine serum albumin (BSA, Sigma-Aldrich) for 30 min at room temperature and incubated with antibody directed against CD13 (#ab33489, Abcam) overnight at 4 °C, followed by PBS washes. Subsequently, Alexa 647-conjugated secondary antibody (#ab150167, Abcam) was introduced and incubated for 1 h at room temperature. Fluorescent images were obtained using an inverted microscope (IX-83, Olympus) and a microscope digital camera (DP80, Olympus). Cross-sectional images were obtained at 2 µm intervals in the vertical direction using a confocal microscopy (FluoView FV1000 confocal, Olympus).

### Statistics and source data

Statistical analyses were done with student t-test. All statistical evaluation was conducted using a one-tailed t-test, assuming independent samples of normal distribution with equal variance. We used Microsoft Excel for calculations and expressed the results as the means ± s.d.

## Acknowledgements

We would like to thank Dr. Misaki Ouchida for graphical assistance, Ms. Harumi Watanabe for administrative assistance, Ms. Yuka Ozaki and Kayo Yano for their technical assistance, and Dr. Peter Karagiannis for reading and editing the paper. This work was supported by the Core Center for iPS Cell Research of Research Center Network for Realization of Regenerative Medicine from the Japan Agency for Medical Research and Development (AMED) [M.K.S.], the Program for Intractable Diseases Research utilizing Disease-specific iPS cells (AMED: 17935423) [M.K.S.], and the Center for Innovation program of Japan Science and Technology Agency (JST) [R.O. and M.K.S.]. R.S. is a recipient of Early Career KAKENHI, iPS Academia Japan, and Sen-shin Medical Research Foundation (SMRF) fellowships. This work was also supported by AMED under Grant No. JP18gm5810008, JSPS KAKENHI Grant No. JP17H02082, and the Kyoto University Hakubi Project.

